# Universal latent axes capturing Parkinson’s patient deep phenotypic variation reveals patients with a high genetic risk for Alzheimer’s disease are more likely to develop a more aggressive form of Parkinson’s

**DOI:** 10.1101/655217

**Authors:** Cynthia Sandor, Stephanie Millin, Andrew Dahl, Michael Lawton, Leon Hubbard, Bobby Bojovic, Marine Peyret-Guzzon, Hannah Matten, Christine Blancher, Nigel Williams, Yoav Ben-Shlomo, Michele T. Hu, Donald G. Grosset, Jonathan Marchini, Caleb Webber

## Abstract

The generation of deeply phenotyped patient cohorts offers an enormous potential to identify disease subtypes but are currently limited by the cohort size and the heterogeneity of the clinical assessments collected across different cohorts. Identifying the universal axes of clinal severity and progression is key to accelerating our understanding of how disease manifests and progresses. These universal axes would accelerate our understanding of how Parkinson’s disease (PD) manifests and progresses through which patients may be appropriately compared appropriately stratified, and personalised therapeutic strategies and treatments can be developed and targeted. We developed a Bayesian multiple phenotype mixed model incorporating the genetic relationships between individuals which is able to reduce a wide-array of different clinical measurements into a smaller number of continuous underlying factors named phenotypic axis. We identify three principal axes of PD patient phenotypic variation which are reproducibly found across three independent, deeply and diversely phenotyped cohorts. Together they explain over 75% of the observed clinical variation and remain robustly captured with a fraction of the clinically-recorded features. The most influential axis was associated with the genetic risk of Alzheimer’s disease (AD) and involves genetic pathways associated with neuroinflammation. Our results suggest PD patients with a high genetic risk for AD are more likely to develop a more aggressive form of PD including, but not limited to, dementia.

## Introduction

A critical challenge in medicine is to understand why the clinical presentations of each patient affected by the same disorder vary. This is especially true for Parkinson’s disease (PD), for which the age of onset, the rate of progression, type and severity of symptoms differ across more than a million people worldwide living with this disease ^1^. To accelerate the identification of disease subtypes, large deeply phenotyped cohorts of PD patients have been created, in which valuable clinical, imaging, biosample and genetic data has been collected, and increasingly with longitudinal monitoring ^2–4^.

Recent studies exploiting these deeply phenotyped cohorts have classified patients into discrete phenotypic subgroups, each displaying a characteristic set of symptoms ^5–7^. To define PD subtypes, most of these studies employ some form of variable selection to create a distance matrix between individuals, followed by clustering methods such as k-means or hierarchical clustering. These methods provide discrete phenotypic groups, which are appealing in their categorical nature but have many shortfalls. Firstly, while selection methods quantify how much variance each phenotype explains, no robust method was used to define a threshold for this measure above which a phenotype contributes to the distance matrix. Consequently, the definition of which phenotypes are essential to group patients and which are irrelevant can be somewhat arbitrary. For example, two recent studies ^5, 8^, using the same Parkinson’s Progression Markers Initiative (PPMI) cohort show divergent results: apathy and hallucinations were key subtype classifiers in the first study ^8^, but not in the second one ^5^, because these variables were not included. Secondly, K-means clustering requires the number of phenotypic groups to be prespecified, and this choice has the potential to be biased towards preconceived expectations with smaller groups ignored or erroneously joined with larger groups. Finally, the creation of discrete groups may not reflect the possibly continuous nature of phenotypic variability and ignores the greater statistical power of continuous traits.

To overcome these limitations, we propose here an approach focused on the continuous variation of phenotypes. Rather than focusing on presence versus absence, or mild versus severe phenotypes, we incorporate the whole spectrum of severity displayed across the population. For this, we applied PHENIX (PHENotype Imputation eXpediated), a multiple phenotype mixed model (MPMM) approach initially developed to impute missing phenotypes ^9^, that can also be exploited for genetically-guided dimensionality reduction of multiple traits. This approach models the phenotypes as a combination of genetic and environmental factors and the genetic component is computed from the correlation matrix between the individual’s genetic data.

Applying PHENIX to the deeply phenotyped UK-based *Discovery* cohort, we identify a small number of axes underlying individual PD patient phenotypic variation that explain the variation in the much larger number of clinically-observed phenotypes. We demonstrate the universality of these axes of phenotypic variation amongst PD patients by independently deriving similar axes in each of three cohorts: UK *Tracking* cohort including 1807 individuals, the UK *Discovery* cohort including 842 PD patients and US PPMI cohort including 439 PD patients that has a different clinical structure from the UK cohorts. We show that this reproducibility is not achieved by other commonly-used dimensionality-reduction methods. Finally, we demonstrate that the most influential axis was associated with the genetic risk of Alzheimer’s disease (AD) suggesting PD patients with a high genetic risk for Alzheimer’s disease are more likely to develop a more aggressive form of PD including dementia symptoms.

## Materials and Methods

### Discovery cohort

We considered 842 PD cases from the *Discovery* cohort constituted of 1700 subjects, including over 1000 people with Parkinson’s, plus 320 healthy controls and 340 individuals thought to be ‘at-risk’ of developing future Parkinson’s. Individuals were required to have at least 90% chance of PD according to UK-Parkinson’s disease brain bank criteria, no alternative diagnosis and disease duration less than 3.5 years. All patients have a clinical assessment repeated every eighteen months and have been already described^4^,^6^. Phenotype data were collected for over a hundred clinical attributes, affecting autonomic, neurological and motor phenotypes (**Supplementary Fig. 1**) and described in the **Supplementary Table 1**. Genotype data were generated using the Illumina HumanCoreExome-12 v1.1 and Illumina InfiniumCoreExome-24 v1.1 SNP arrays.

### UK Tracking Parkinson’s study

We considered 1807 PD cases from the *Tracking* Parkinson’s cohort, which was already described in detail by Malek *et al*. ^*2*^ and was used to identify the impact of mutations within glucocerebrosidase gene (*GBA*) on different PD clinicals manifestations ^10^. Genotype data were generated using the Illumina Human Core Exome array.

### PPMI cohort

The PPMI cohort (http://www.ppmi-info.org) was already described in detail (including PPMI protocol of recruitment and informed consent) by Marrek *et al*. ^11^. We downloaded data from the PPMI database on September 2017 in compliance with the PPMI Data Use Agreement. We considered 472 newly-diagnosed typical PD subjects: subjects with a diagnosis of PD for two years or less and who are not taking PD medications. We used the baseline (t=0) of clinical assessments, described in detail in the **Supplementary Table 2**. We excluded any individual with > 5% of missing data (437 individuals included). Participants have been genotyped using two genotyping arrays, ImmunoChip ^12^ and NeuroX ^13, 14^. As more participants were genotyped on NeuroX array, we used the genotype data of the NeuroX chip.

## Methods

### Genotype: quality control & Imputation

Quality control was carried out independently using PLINK v1.9 ^15^ (SI). Imputation of unobserved and missing variants was carried out separately for each cohort (SI)

### Phenotypic axis

Our continuous measures of severity are based on a multiple phenotypes mixed model approach (MPMM) named PHENIX (PHENotype Imputation eXpediated) which includes genetic relationships between individuals, and is designed to impute missing phenotypes ^9^. To impute missing phenotypes, PHENIX reduces the variation within a cohort to a smaller number of underlying factors that are then used to predict individual missing values. Here, we exploit the identification of these underlying factors as providing the latent axes of patient variation which underlie a larger number of clinically observed phenotypes. The outcome is that the many clinical phenotypes (sometimes missing for some individuals) of each individual are represented through a smaller number of underlying latent variables of phenotypic variation that manifest the observed clinical phenotypes, which we name herein as *phenotypic axes*.

PHENIX ^9^ use a Bayesian multiple-phenotype mixed model (MPMM), where the correlations between clinical phenotypes (Y) are decomposed into a genetic and a residual component with the following model: Y=U+e, where U represents the aggregate genetic contribution (whole genotype) to phenotypic variance and e is idiosyncratic noise. As the estimation of maximum likelihood covariance estimates can become computationally expensive with increasing number of phenotypes, PHENIX uses a Bayesian low-rank matrix factorization model for the genetic term U such as: U = Sβ, in which β is can be used to estimate the genetic covariance matrix between phenotypes and S represents a matrix of latent components that each follow ∼N (0,G) where G is the Estimate of Relatedness Matrix from genotypes. The resulting latent traits (S) are used as phenotypic axes, each representing the severity of a number of non-independent clinical phenotypes. The details to run PHENIX and extract the phenotypic axes are given in the **Supplemental Information**.

### Disease phenotypic axis

We derived disease phenotypic axis consisting to replace the general genetic component in a MPMM by a disease risk genetics component. To calculate a disease relatedness matrix, we considered only genetic variants (after pruning) associated with human complex traits. For different complex human traits with GWA results publically available (**Supplementary Table 01)**, we calculated a disease relatedness, that we used subsequently to derive phenotypic axes (**Supplementary Information**).

## Results

### Three continuous measures capture 75% of the clinical variation

Initially, we generated phenotypic axes from a cohort of 842 PD patients (*Discovery* cohort) which had been genotyped and phenotypically characterised with 40 clinical assessments (**Supplementary Table 1**). Each latent axis reflected a number of co-varying observed clinical assessments. Among the phenotypic axes that explained more than 5%, Axes 1, 2 and 3 explained 39.6%, 28.7% and 6.8% of the clinical variation respectively. Together, these 3 top axes account for over 75% of the clinically-observed variation (**Supplementary Fig. 2**). To examine whether similar phenotypic axes are obtained in different deeply phenotyped PD cohorts, we derived phenotypic axes within an independent cohort of 1807 PD individuals from the UK *Tracking* cohort ^2^ that had made similar clinical observations to the *Discovery* cohort. We found significant Pearson’s correlation coefficients between each cohort’s first three phenotypic axes: Axis 1 r=0.92 (p=3 × 10^−13^), Axis 2 r=0.89 (p=4 × 10^−11^), Axis 3 r=0.72 (p=5 × 10^−6^) (**Fig. 1**). Nevertheless, a major concern was that the identification of the same phenotypic axes might, at least in part, be due to the very similar structure of the clinical phenotyping between the two UK cohorts. To address this, we examined the independent US-based PPMI cohort consisting of 439 sporadic PD individuals that had been clinically phenotyped following a substantially different protocol to the UK cohorts. After deriving phenotypic axes in the PPMI cohort, we found significant similarities between the first three phenotypic axes derived for both the *Discovery*-UK and PPMI-US cohorts: the coefficients of determination (R^2) between three first axes across different categories of clinical phenotypes from each cohort were: Axis1: 0.665 (p=0.048), Axis 2: 0.914 (p=0.003) and Axis 3: 0.754 (p=0.025) (**Fig. 2 & Supplementary Figure 3**). These consistent similarities in the axes of phenotypic variation independently derived for each of three different PD cohorts demonstrates the reproducibility of these axes of phenotypic variation amongst Parkinson’s patients. Finally, by comparing PHENIX with other methods of dimensionality reduction, specifically Principle Component Analyses (PCA), Multidimensional Scaling (MDS) and Independent component analysis (ICA), only the dimensions discovered by the MPMM model, PHENIX, were significantly correlated between both cohorts and thus no other method was able to identify similar axes of phenotypic variation across UK and US PD cohorts (**Fig. 2**).

**Fig. 1.**
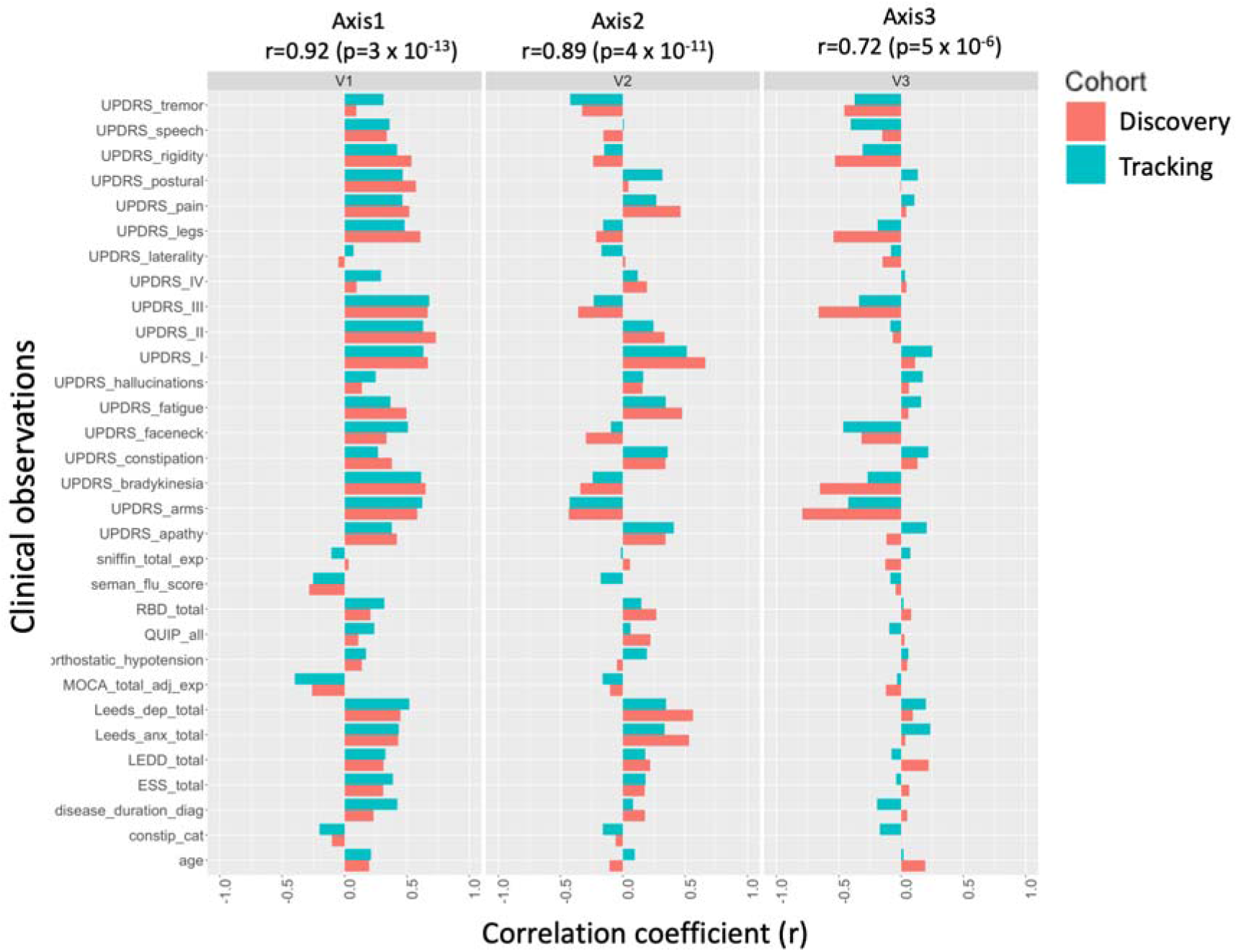
The clinical phenotypes of two independent deeply phenotyped Parkinson’s disease cohorts identify the same phenotypic axes. Results were consistent in two independents cohorts (842 *Discovery* and 1807 *Tracking* patients). Examination of these two separate Parkinson’s disease cohorts, using independent derivation of the phenotypic axes in each, showed significant correlations between each cohort’s first three axes. Correlations between the axes from each cohort are Axis 1 r=0.92 (p=3 x 10-13), Axis 2 r=0.89 (p=4 x 10-11), Axis 3 r=0.72 (p=5 x 10-6). The correlation coefficient (x-axis) between each axis derived in each cohort (blue: *Discovery* vs red: *Tracking*) and each clinical observation (y-axis) is shown.

**Fig. 2.**
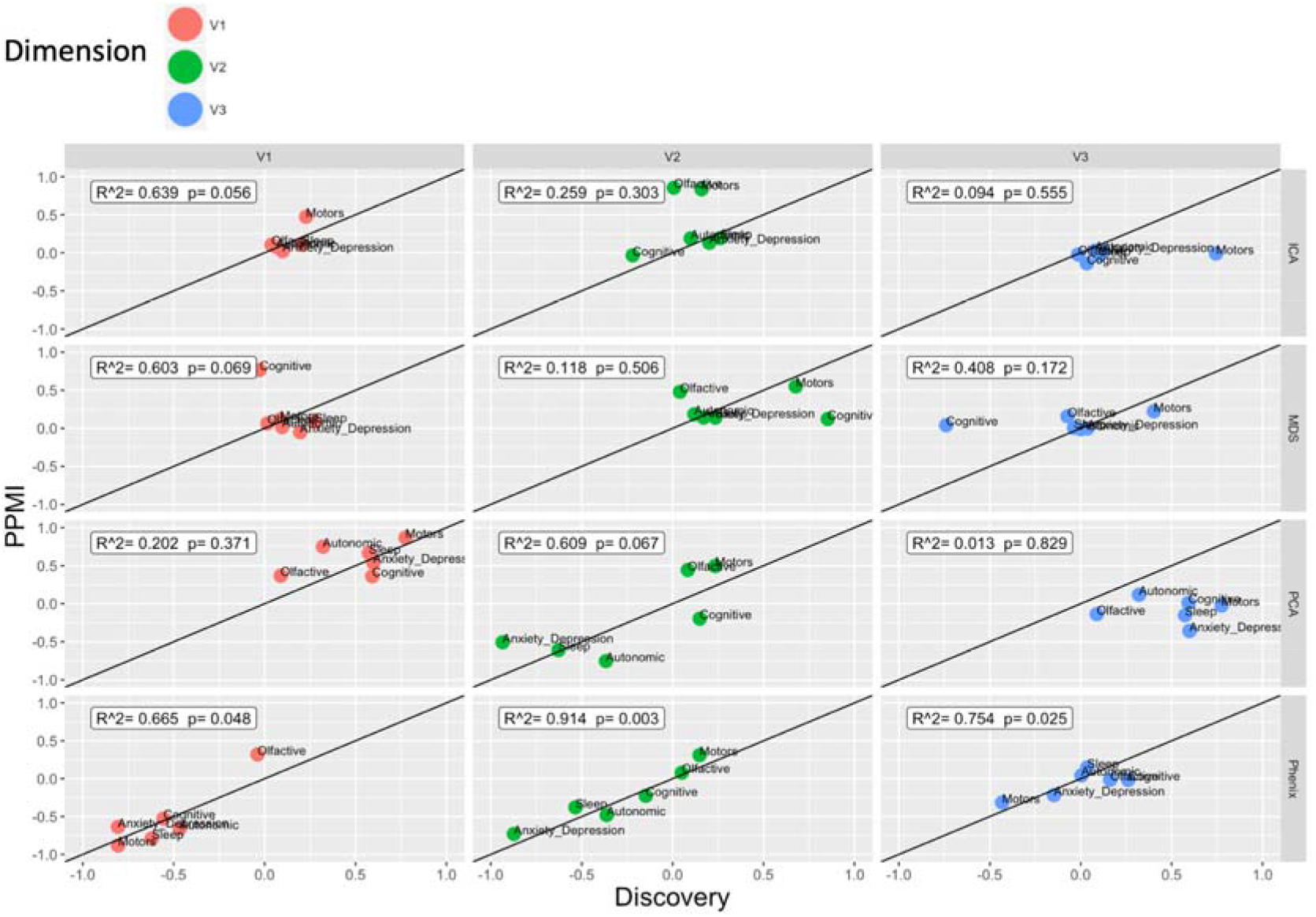
The reduced dimensions in other dimensionality reduction methods fail to align between differently but deeply phenotyped UK and US Parkinson’s disease cohorts. We compared the ability of different dimensionality reduction methods (independent component analysis (ICA), Multidimensional scaling (MDS), Principal component analysis (PCA) and phenotypic axis based on the PHENIX multiple phenotype mixed model) to phenotypically align two deeply phenotyped Parkinson’s disease cohorts, specifically the *Discovery* (842 individuals) and PPMI (439 sporadic Parkinson’s disease) cohorts. The x-axis and y-axis represent the correlation coefficient between each continuous variable with clinical observation associated with a specific symptom category in *Discovery* and PPMI cohort respectively. Each column panel and colour of points (“Axis”) represents the dimension level of each underlying dimension. All points on the diagonal would represent a perfect phenotypic alignment of both cohorts. We examined the relationship between correlation derived from both cohorts by performing a linear regression: R^2 and p correspond to the coefficient of determination and the p-value respectively. Only the dimensions discovered by the MPMM model, PHENIX, show a significant relationship between both cohorts: MPMM phenotypic axes (R^2^=0.86, p=2×10-8), MDS (R^2^=0.11, p=0.18), ICA (R^2^=0.17, p=0.16) and PCA (R^2^=0.31, p=0.06).

### Each phenotypic axis represents a distinct set of clinical features

To interpret the clinical relevance of each phenotypic axis, we examined the correlation between individual clinical features and the phenotypic axes (**Table 1 & Fig. 1 & Supplementary Figure 4**). We observed that each phenotypic axis corresponded to a subset of clinical features, differing in both extents and directions of severity. Axis 1 represented worsening non-tremor motor phenotypes, anxiety and depression accompanied by a decline of the cognitive function (**Table 1 & Fig. 3**). Worsening anxiety and depression were also features of Axis 2, in addition to increasing severity of autonomic symptoms and increasing motor dysfunction. Axis 3 was associated with general motor symptom severity including rigidity, bradykinesia and tremor of the whole body independently of non-motor features. The contribution of different phenotypes to these axes was therefore highly variable. Specific aspects of motor dysfunction were important factors in defining the majority of axes. Anxiety and depression were also relatively important features, but only for axes explaining the largest amounts of variation. Conversely, cognitive impairment was associated only with Axis one. However, this observation must be weighted by the fact that cognitive impairment/dementia are reported at a later disease stage and thus likely under-represented in recently diagnosed cases.

**Fig. 3.**
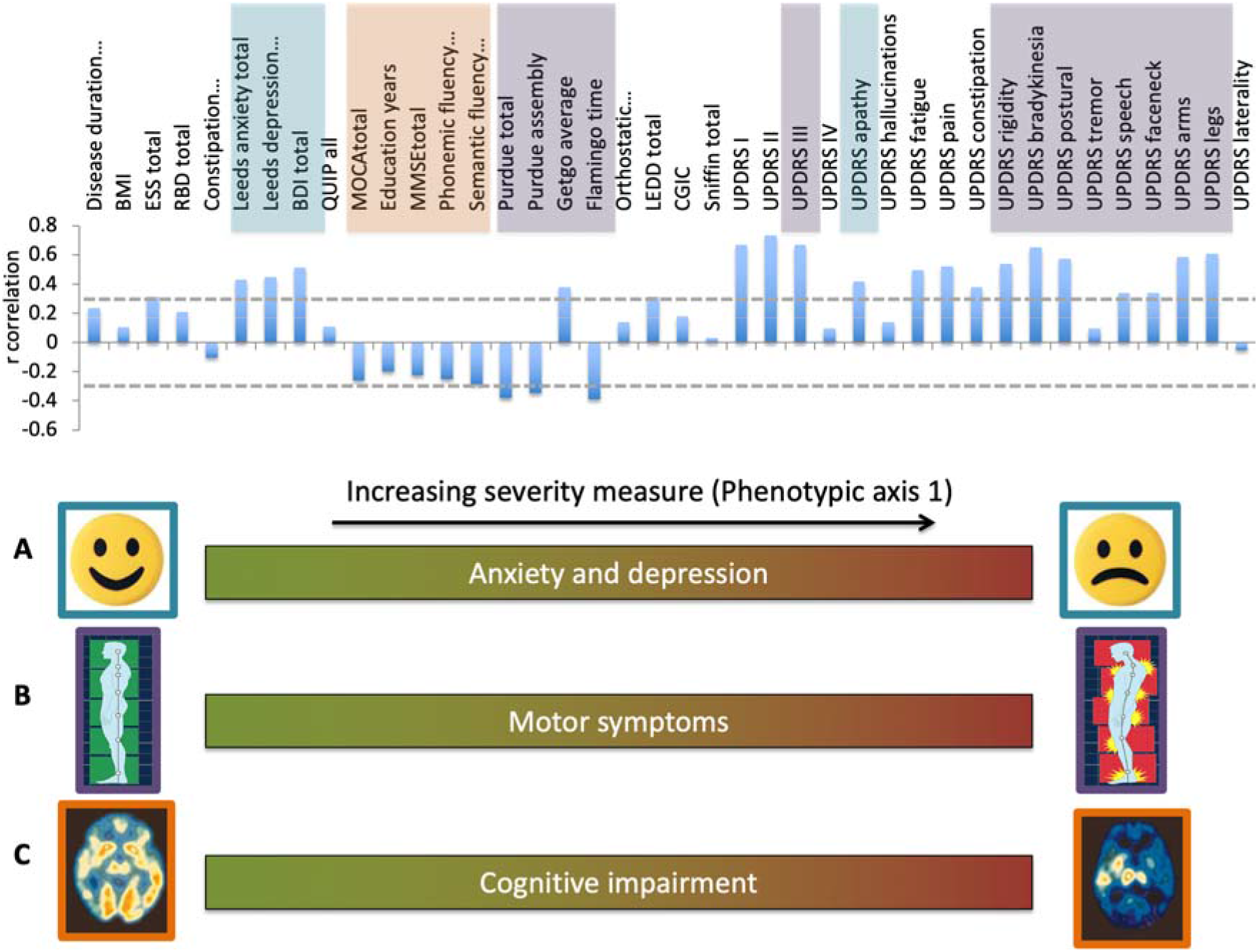
The correlation of individual clinically-measured Parkinson’s disease phenotypes with an underlying Phenotypic Axis 1. Modelling patient clinical phenotypes as a combination of genetic and environmental factors revealed three phenotypic severity axes (**Fig**.**1**), each representing a continuous pattern of variation between multiple co-varying clinical phenotypes. In Axis 1 (shown), (A) clinical measures relating to anxiety and depression and apathy are significantly and positively correlated with an individual’s score along this axis; patients with a higher axis score have more severe mood and neuropsychiatric problems. (B) The severity of motor phenotypes is positively correlated with this phenotypic axis; patients with a higher axis score is associated with more severe motor phenotypes (C) Cognitive tests were negatively correlated with this component (the patients that score high in these cognitive tests have less cognitive impairments); individuals with a high score for this component suffer from more severe anxiety, depression and displayed more cognitive impairment and motor symptoms.

**Table 1:**
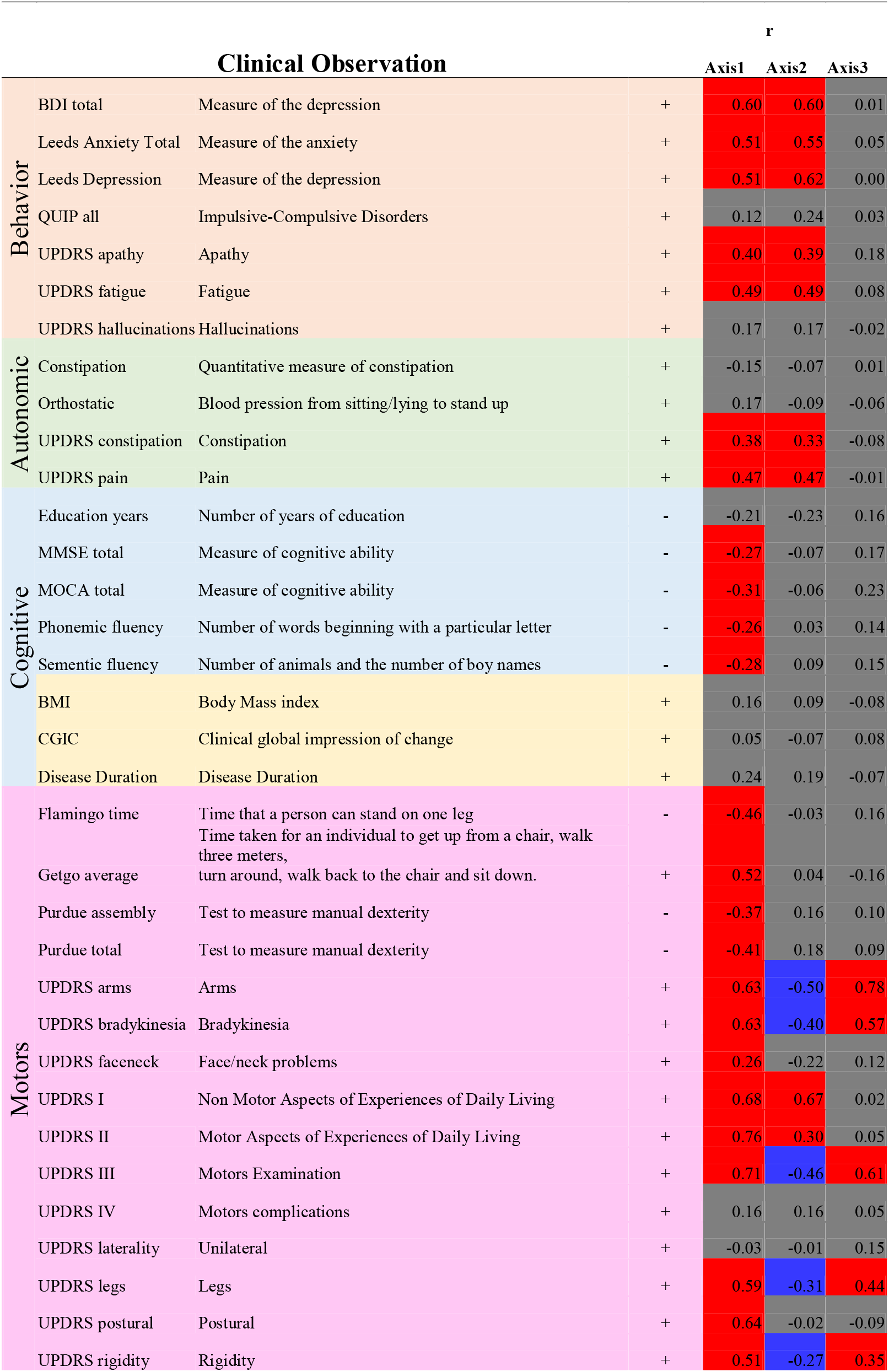
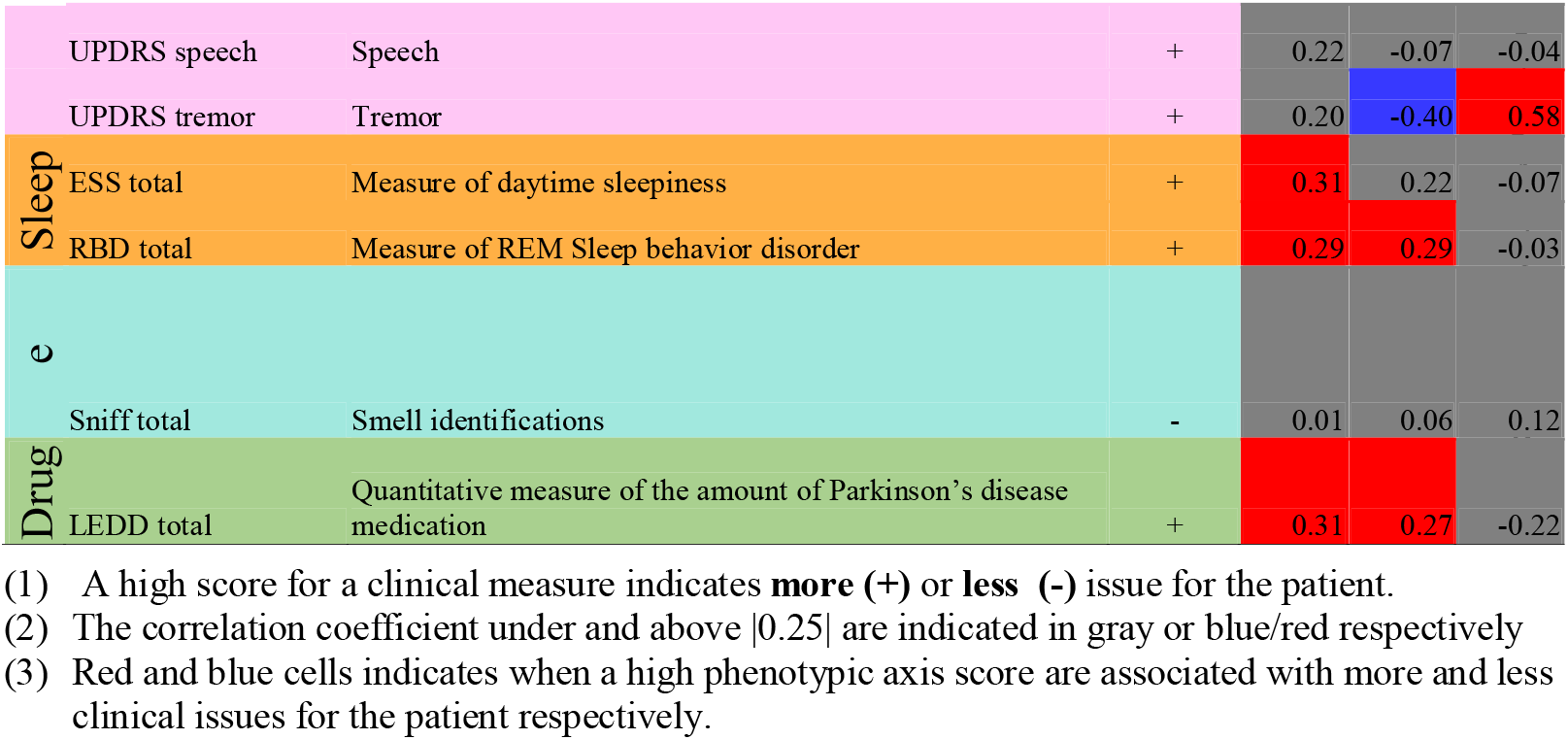
Correlation between each axis and each clinical phenotypic measure

Although each phenotypic axis is associated with a distinct set of clinical features, they are not independent but instead strongly correlated (**Supplementary Figure 5**). We find no significant relation between the phenotypic axes and principal components of genetic ancestry (**Methods**) suggesting that the phenotypic axes are not biased by the population structure (**Supplementary Figure 5, Supplementary Table 3**). However, as previously reported, gender influences clinical symptoms ^4^ and we also observe a significant association between gender and Axis 2 (**Supplementary Table 3**, p=4.5×10^−5^).

To assess to what extent the phenotypic axes might be affected by the number of clinical observations, within the *Discovery* cohort we compared the phenotypic axes built on all clinical features with phenotypic axes generated with incomplete sets of randomly-selected clinical features. We observed a strong correlation (r > 0.8) between each of the two first phenotypic axes built with as few as 50% of the clinical variables and their respective original phenotypic axes, suggesting that these two axes are extremely robust in terms of the numbers of clinical variables considered (**Supplementary Figure 6**).

### The integration of genetic relationships between patients improves capture of the Parkinson’s disease clinical variation and reproducibility

The PHENIX MPMM approach employed here to derive phenotypic axes exploits the genetic relatedness between individuals derived from genotypic similarity to further decompose random effects into kinship effects between individuals. In its original application to imputing missing phenotypes, PHENIX outperforms other imputation approaches when the heritability (h^2^) of a phenotype increased ^9^. Similarly, when randomly removing and re-imputing 10% of observed data, the quality of the imputation of PD clinical assessments was in general better when considering the genetic relatedness between individuals as compared to excluding this information (**Supplementary Figure 7**), suggesting that the resulting phenotypic axes better capture PD heterogeneity when including genetic information. Moreover, we found a higher agreement between the phenotypic axes derived by integrating the genetic relationship between patients of different cohorts than when the phenotypic axes were derived ignoring the genetic relationships (**Supplementary Figure 8**). Specifically, the coefficient of determination reflecting the agreement between the axes derived from the Discovery and those derived from the PPMI cohorts were from Axis 1 to 3: 0.665 (p=0.048), 0.914 (p=0.003) and 0.754 (p=0.025) when including the genetic similarity between patients as compared to 0.604 (p=0.069), 0.908 (p=0.003) and 0.001 (p=0.991) without. Together, these findings demonstrate that the integration of genetic relationship between patients enhances the resulting phenotypic axes’ ability to reproducibly capture PD clinical variation.

### Metanalysis of Genome Wide Association Studies with phenotypic axes as unique and universal quantitative traits

Each phenotypic axis provides a quantitative trait enabling the genetics underlying patient variation to be studied by performing a Genome Wide Association Study (GWAS) via a regression model with the covariates age, gender, and two genetic principal components (to account for any underlying population substructure) in each individual cohort. As three phenotypic axes were similar across each individual cohort (*Discovery, Tracking* and PPMI) and to increase statistical power to detect an significant association, we conducted a meta-analysis of each phenotypic axis genome-wide association studies using a common set of 4211937 variants across 3088 individuals. A significant departure from the expected quantiles was observed for Axis 1 (meta-analysis combining the summary statistic of three individual GWAS [*Discovery*-Tracking-PPMI]) (**Supplementary Figure 9**), but no variant surpassed genome-wide significance (**Supplementary Figure 10**). Although we did not observe a significant genome-wide association, the use of universal phenotypic axes significantly unable us to conduct meta-analysis and thus to increase the statistical power to identify genetic variants through their ability to align differently deeply phenotyped cohorts and reduce the number of traits tested.

Next, we re-examined genetic associations for each of the three phenotypic axes for three major PD risk genes, namely *SNCA, GBA* and *LRRK2*. We found a indicative local association signal but however un-significant at GWA level with Phenotypic Axis 1 for a variant in *SNCA*: 4: 90758437 (p-value=1.7×10^−4^, **Supplementary Figure 11A**) which is in high LD with rs1348224 (r^2^ > 0.8), a SNP previously associated with PD with dementia and dementia with Lewy bodies ^20^. SNP rs1348224:G allele (minor allele) had a negative effect on Phenotypic Axis 1, thus a protective effect for cognitive impairment, which is consistent with a protective effect for PD with dementia and dementia with Lewy bodies previously reported for this locus ^20^. We also found a indicative local association signal (p-value=1.1×10^−4^) with Phenotypic Axis 3 for an intronic variant in *LRRK2* (**Supplementary Figure 11B)**. Both *SNCA* and *LRRK2* variants were each nominally associated with only one phenotypic axis (**Supplementary Table 4**), suggesting distinct pathogenic mechanisms.

### Parkinson patients carrying a high genetic risk for Alzheimer’s are more likely to develop a more aggressive form of Parkinson’s

To better understand the genetics risk factors influencing the phenotypic axis, we calculated a disease-risk relatedness matrix in the MPMM, based on genetic variants associated with different complex human traits. For example, replacing the overall genetic similarity by how similar people are in their risk of diabetes or depression. By examining the proportion of phenotypic variation explained by different phenotypic axis derived using these different disease risks as compared to the original phenotypic axes derived using the entire genotype, we showed that the phenotypic axis derived using Alzheimer’s disease (AD) genetic risk significantly outperforms (capture more patient phenotypic variation) the original phenotypic axes (**Fig. 4)**

**Fig. 4.**
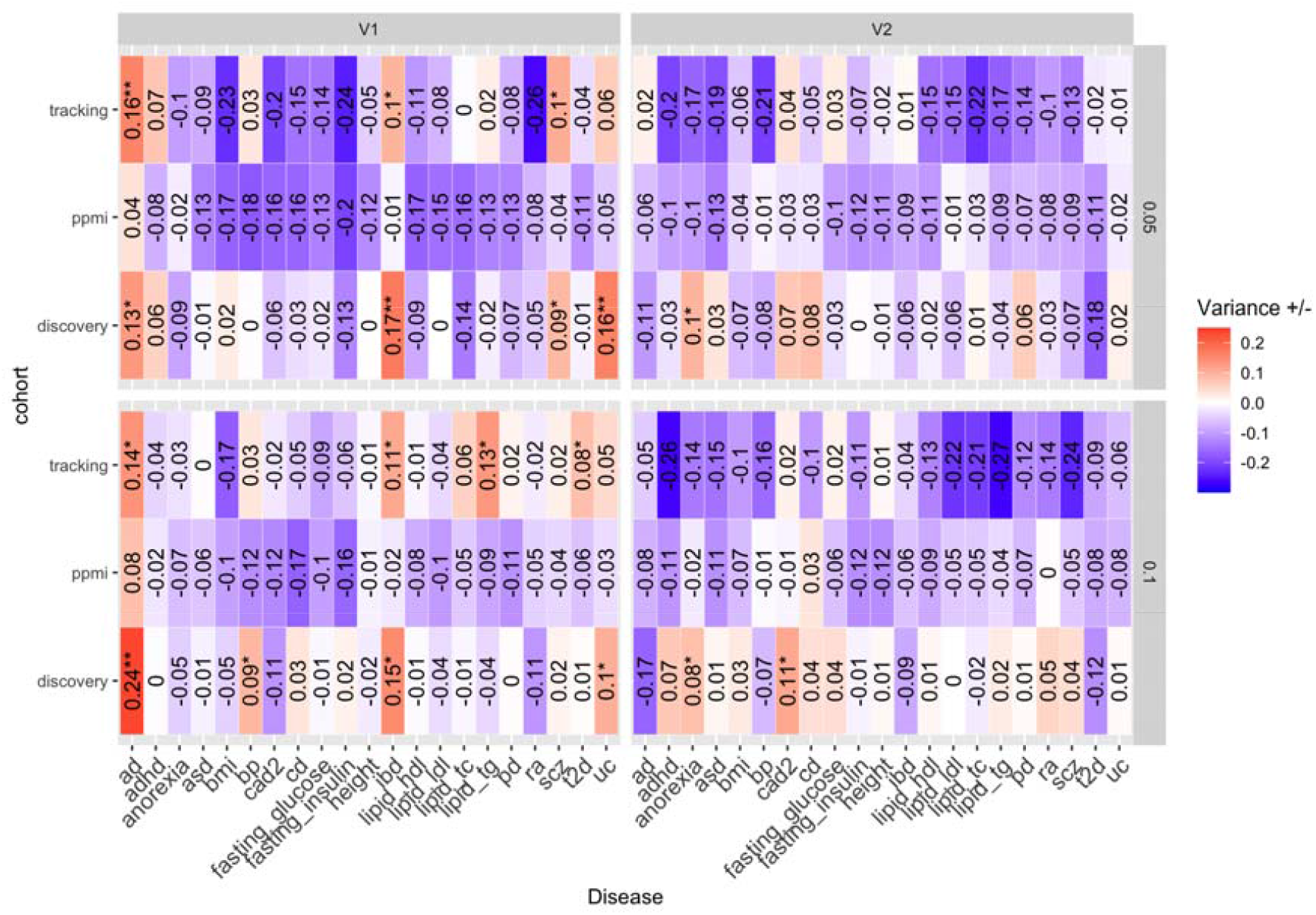
Alzheimer’s disease phenotypic axis significantly outperforms the original phenotypic axis. These heatmap plots represents for two first phenotypic axes (left=V1 and right=V2), in the 3 cohorts (row1=Discovery, row2=Tracking & row 3=) the excess(red) or deficit of the phenotypic variance explained by the different disease axes (columns) compared with the original phenotypic axes.

This result proposes that PD patients carrying a high genetic risk for AD are more likely to develop a more aggressive form of PD including dementia symptoms: Axis 1 represents worsening non-tremor motor phenotypes, anxiety and depression accompanied by a decline of the cognitive function (preprint Table 1 & Fig. 2) Testing this hypothesis in the PPMI cohort, we found a significant relationship between Phenotypic Axis 1 and CSF Aβ1-42, a biomarker strongly associated with future conversion to dementia. Secondly, as for AD genetics risk and unlikely to PD genetics risk, we found an association of phenotypic axis 1 risk variants with microglia-expressed genes, in both the SN and the cortex suggesting that the neuro-inflammation play a key role in the development a more aggressive form of PD, but not in the PD onset-risk. Finally we observed the phenotypic axis one is associated with rapid progression of multiple clinical symptoms suggesting that AD genetic risk score in PD patients could be used as a predictor of progression

## Discussion

We propose here a novel approach to quantifying diverse patient phenotypes on a continuous scale via the use of phenotype axes. This approach overcomes many of the limitations associated with the clustering methods previously used to classify PD heterogeneity. By applying our approach to three independent and deeply phenotyped cohorts, we demonstrate the universality of these axes of phenotypic variation amongst PD patients. We also showed that our axes are robustly derived when reducing the number of clinical features considered and, unlike other dimensionality reduction methods, the PHENIX MPMM approach is the only method tested here that is able to identify the same phenotypic axes underlying PD patient variation between individuals from different cohorts. The phenotypic axes have multiple applications in PD precision medicine. We found that PD patients carry on a high genetic risk load for Alzheimer’s disease can develop a more clinical aggressive PD form including dementia symptoms.

Our approach was able to identify representative quantitative variables that are clinically relevant to previously-defined categorical PD subtypes. A number of known comorbidities were represented among the phenotype axes. Anxiety and depression are highly correlated in PD patients, both of which are correlated with Axes 1 and 2 ^26^. Rigidity and bradykinesia are also linked, possibly due to shared physiology ^27^, and varied in the same direction along Axis 3. Lawton *et al*. reported five PD subgroups, by using the same *Discovery* cohort but following a k-means clustering approach ^6^. We examined the distribution of phenotypic axis score across these five PD subgroups (**Supplementary Figure 16**) and noted that the 5^th^ subgroup of patients, characterised by severe motor, non-motor and cognitive disease, with poor psychological well-being clinical symptoms, were systematically associated with high severity score for all three of our phenotypic axes. Inversely, the first PD subgroup characterised by mild motor and non-motor disease (group affected by fewer clinical symptoms) displayed a low severity score for our three phenotypic axes. Furthermore, we observed that the individuals of subgroups 4 and 5, characterised by poor psychological well-being, had high severity scores for phenotypic axis 2, the axis most associated with depression and anxiety symptoms. These observations demonstrate some consistency between subgroups defined with k-means and our phenotypic axis severity score. The agreement of these phenotype axes with previously observed correlations provides further support for underlying biological themes, but their reinterpretation as robust continuous traits likely provides a better approximation of how the underlying biology contributes, as opposed to a cut-off off for a phenotype. Specifically, the unimodal character of the phenotypic axis distributions (**Supplementary Figure 17**) suggests here that the development of continuous measures is more appropriate than clustering according to an arbitrary threshold.

The phenotypic axes identified were robust in terms of the number of clinical features considered and enable the alignment of patients from different cohorts with different clinical phenotyping structures. The corollary is that Phenix did not require the variable selection common in PD clustering approaches, and it can also guide clinicians in determining which clinical assessments are essential to capture PD heterogeneity. Deep phenotyping is burdensome to both patient and clinician and many of the measures exploited here are compound scores summarising aspects of functioning. Further work identifying the minimally burdensome observations that enable robust scoring of patients along these phenotypic axes would facilitate their utility and adoption across the PD clinical community, bringing increased power to the discovery of influencing factors. Finally, the MPMM approach can be readily extended to include longitudinal data to determine the phenotypic axes associated with disease progression while simultaneously dealing with missing data, which is a common problem in longitudinal studies.

In conclusion, these universal axes have the potential to accelerate our understanding of how PD presents in individual patients, providing more robust and objective quantitative traits through which patients may be appropriately compared, through which the underlying disease-modifying mechanism can be understood and appropriately stratified/personalised therapeutic strategies and treatments can be developed.

## Acknowledgments

The work was supported by the Monument Trust Discovery Award from Parkinson’s UK. Oxford Genomics Centre at the Wellcome Centre for Human Genetics, Oxford is Funded by Wellcome Trust (grant reference 090532/Z/09/Z and MRC Hub grant G0900747 91070) Samples and associated clinical data were supplied by the Oxford Parkinson’s Disease Centre (OPDC) study, funded by the Monument Trust Discovery Award from Parkinson’s UK, a charity registered in England and Wales (2581970) and in Scotland (SC037554), with the support of the National Institute for Health Research (NIHR) Oxford Biomedical Research Centre based at Oxford University Hospitals NHS Trust and University of Oxford, and the NIHR Comprehensive Local Research Network. CW is supported by a UK DRI fellowship funded by Medical Research Council (MRC), Alzheimer’s Society and Alzheimer’s Research UK. CW and CS are supported by Computational Science Program funded by Michael J. Fox Foundation. JM acknowledges funding for this work from the European Research Council (ERC; grant 617306). We thank the Oxford Genomics Centre at the Wellcome Centre for Human Genetics, Oxford) for the generation genotyping data.

